# Disentangled behavioral representations

**DOI:** 10.1101/658252

**Authors:** Amir Dezfouli, Hassan Ashtiani, Omar Ghattas, Richard Nock, Peter Dayan, Cheng Soon Ong

## Abstract

Individual characteristics in human decision-making are often quantified by fitting a parametric cognitive model to subjects’ behavior and then studying differences between them in the associated parameter space. However, these models often fit behavior more poorly than recurrent neural networks (RNNs), which are more flexible and make fewer assumptions about the underlying decision-making processes. Unfortunately, the parameter and latent activity spaces of RNNs are generally high-dimensional and uninterpretable, making it hard to use them to study individual differences. Here, we show how to benefit from the flexibility of RNNs while representing individual differences in a low-dimensional and interpretable space. To achieve this, we propose a novel end-to-end learning framework in which an encoder is trained to map the behavior of subjects into a low-dimensional latent space. These low-dimensional representations are used to generate the parameters of individual RNNs corresponding to the decision-making process of each subject. We introduce terms into the loss function that ensure that the latent dimensions are informative and disentangled, i.e., encouraged to have distinct effects on behavior. This allows them to align with separate facets of individual differences. We illustrate the performance of our framework on synthetic data as well as a dataset including the behavior of patients with psychiatric disorders.

## 1 Introduction

There is substantial commonality among humans (and other animals) in the way that they learn from experience in order to make decisions. However, there is often also considerable variability in the choices of different subjects in the same task [Carroll and Maxwell, 1979]. Such variability is rooted in the structure of the underlying processes; for example, subjects can differ in their tendencies to explore new actions [e.g., Frank et al., 2009] or in the weights they give to past experiences [e.g., den Ouden et al., 2013]. If meaningfully disentangled, these factors would crisply characterise the decision-making processes of the subjects, and would provide a low-dimensional latent space that could be used for many other tasks including studying the behavioral heterogeneity of subjects endowed with the same psychiatric labels. However, extracting such representations from behavioral data is challenging, as choices emerge from a complex set of interactions between latent variables and past experiences, making disentanglement difficult.

One promising approach proposed for learning low-dimensional representations of behavioral data is through the use of cognitive modelling [e.g., Navarro et al., 2006, Busemeyer and Stout, 2002]; for example using a reinforcement learning framework [e.g., Daw, 2011]. In this approach, a parametrised computational model is assumed to underlie the decision-making process, and the parameters of this model – such as the tendency to explore and the learning rate – are found by fitting each subject’s choices. Their individual parameters are treated as the latent representations of each subject. This approach has been successful at identifying differences between subjects in various conditions [e.g., Yechiam et al., 2005]; however, it is constrained by its reliance on the availability of a suitable parametric model that can capture behavior and behavioral variability across subjects. In practice, this often leads to manually designing and comparing a limited number of alternative models which may not fit the behavioral data closely.

An alternative class of computational models of human decision-making involves Recurrent Neural Networks (RNNs) [e.g., Dezfouli et al., 2019, 2018, Yang et al., 2019]. RNNs can model a wide range of dynamical systems [Siegelmann and Sontag, 1995] including human decision-making processes, and make fewer restrictive assumptions about the underlying dynamical processes. However, unlike cognitive models, RNNs typically: (i) have large numbers of parameters, which (ii) are hard to interpret. This renders RNNs impractical for studying and modelling individual differences.

Here we develop a novel approach which benefits from the flexibility of RNNs, while representing individual differences in a low-dimensional and interpretable space. For the former, we use an autoencoder framework [Rumelhart et al., 1985, Tolstikhin et al., 2017] in which we take the behaviors of a set of subjects as input and automatically build a low-dimensional latent space which quantifies aspects of individual differences along different latent dimensions (Figure 1). As in an hyper-networks [Ha et al., 2016, Karaletsos et al., 2018], the coordinates of each subject within this latent space are then used to generate the parameters of an RNN which models the decision-making processes of that subject. To address interpretability, we introduce a novel contribution to the autoencoder’s loss function which encourages the different dimensions of the latent space to have separate effects on predicted behavior. This allows them to be interpreted independently. We show that this model is able to learn and extract low-dimensional representations from synthetically-generated behavioral data in which we know the ground truth, and we then apply it to experimental data.

**Figure 1:**
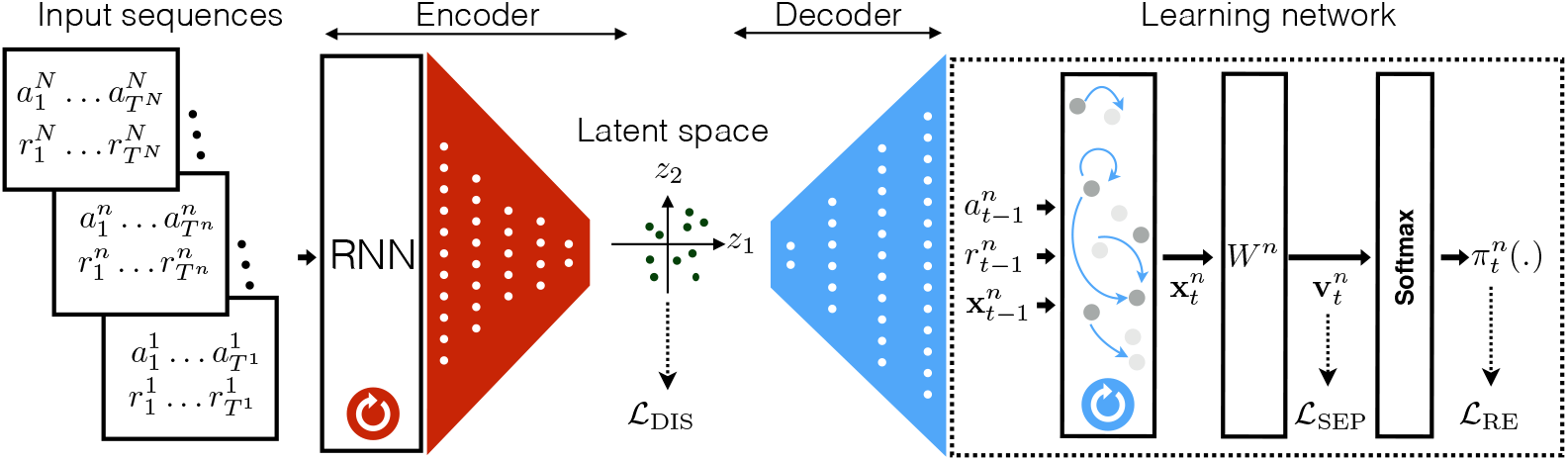
The model comprises an encoder (shown in red) and decoder (blue). The encoder is the composition of an RNN and a series of fully-connected layers. The encoder maps each whole input sequence (in the rectangles on the left) into a point in the latent space (depicted by (*z*_1_, *z*_2_)-coordinates in the middle) based on the final state of the RNN. The latent representation for each input sequence is in turn fed into the decoder (shown in blue). The decoder generates the weights of an RNN (called the learning network here) and is shown by the dotted lines on the right side. The learning network is the reconstruction of the closed-loop dynamical process of the subject which generated the corresponding input sequence based on experience. This takes as inputs the previous reward, 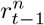, previous action, 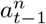, and its previous state, 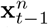, and outputs its next state 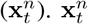 is then multiplied by matrix *W^n^* to generate unnormalised probabilities (logits) 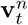 for taking each action in the next trial, which are converted (through a softmax) to actual probabilities, 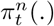. The negative-log-likelihoods of the true actions are used to define the reconstruction loss, 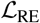, which along with 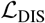 and 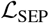 are used to train the model. The 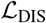 term induces disentanglement at the group level, and 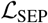 separates the effects of each dimension of the latent space on the output of the learning network.

## 2 The model

### Data

The data 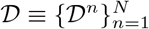 comprise *N* input sequences, in which each sequence *n* ∈ {1…*N*} consists of the choices of a subject on a sequential decision-making task. In input sequence *n* ∈ {1,…, *N*} the subject performs *T^n^* trials, 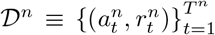, with action 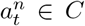 on trial *t* chosen from a set 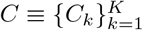, and reward 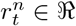.

### Encoder and decoder

We treat decision-making as a (partly stochastic) dynamical system that maps past experiences as input into outputs in the form of actions. As such, the dataset 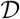 may be considered to contain samples from the output of potentially *N* different dynamical systems. The aim is then to turn each sequence 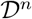 into a vector, **z***^n^*, in a latent space, in such a way that **z***^n^* captures the characteristic properties of the corresponding dynamical system (whilst avoiding over-fitting the particular choices). In this respect, the task is to find a low-dimensional representation for each of the dynamical systems in an unsupervised manner. We take an *autoencoder-inspired* model to achieve this, in which an encoder network is trained to process an entire input sequence into a vector in the latent space. A decoder network then takes as input this vector and recovers an approximation to the original dynamical system. Along with additional factors that we describe below, the model is trained by minimising a reconstruction loss which measures how well the generated dynamical system can predict the observed sequence of actions. The latent representation is thus supposed to capture the “essence” of the input sequence. This is because the latent space is low-dimensional compared to the original sequence and acts as an information bottleneck; as such the encoder has to learn to encode the most informative aspects of each sequence.

The model architecture is presented in Figure 1. The first part is an encoder RNN (shown in red), and is responsible for extracting the characteristic properties of the input sequence. It takes the *whole* sequence as input and outputs its terminal state. This state is mapped into the latent space through a series of fully-connected feed-forward layers. The encoder is:

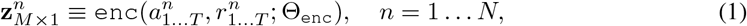

in which Θ_enc_ are the weights of the RNN and feed-forward layers, and *M* is the dimensionality of the latent space (see Supplementary Material for the details of the architecture).

The second part of the model is a feed-forward decoder network (shown in blue) with weights Θ_dec_, which takes the latent representation as input, and outputs a vector Φ*^n^*,

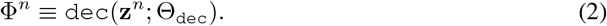

As in a hyper-network, vector Φ*^n^* contains the weights of a second RNN called the *learning network*, itself inspired by [e.g., Dezfouli et al., 2019, 2018], and described below. The learning network apes the process of decision-making, taking past actions and rewards as input, and returning predictions of the probability of the next action. Making Φ*^n^* such that the learning network reconstructs the original sequence is what forces **z***^n^* to encode the characteristics of each subject.

### Learning network

The learning network is based on the Gated Recurrent Unit architecture [GRU; Cho et al., 2014] with *N_c_* cells. This realizes a function *f^n^*, which at time-step *t* maps the previous state 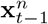, action 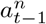 and reward 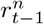, into a next state,

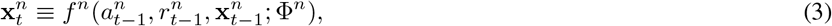

with predictions for the probabilities of actions arising from a weight matrix 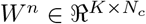

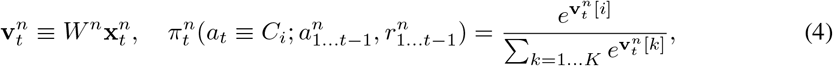

where 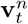 represent the logit scores for each action (unnormalised probabilities), and *π_t_*(.) are the probabilities of taking each action at time *t*. 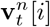 represents the ith element of 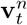. For *N_c_* GRU cells and *K* actions, function *f^n^* requires 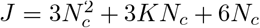 parameters. Φ*^n^* consists of these plus the *KN_c_* parameters of *W^n^*. Note that this RNN, which models humans decision-making processes, should not be confused with the RNN in the encoder, which extracts and maps each input sequence into the latent space.

### Summary

The encoder network takes an input sequence and maps it into its latent representation. The decoder network then takes the latent representation and generates the weights of an RNN that is able to predict the actions taken in the input sequence. The next section describes in detail how the network weights Θ_enc_ and Θ_dec_ are learned end-to-end.

## 3 Training objective

The training loss function has three components: (i) a reconstruction loss which penalizes discrepancies between the predicted and actual input sequence, (ii) a group-level disentanglement loss which encourages sequences to spread independently across the dimensions of the input sequence, (iii) a separation loss which favors dimensions of the latent space that have separate effects on the behavior generated by the learning networks.

### Reconstruction loss

Each sequence 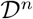 in 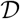 is passed through the encoder to obtain the corresponding latent representation. The latent representation is then passed through the decoder to generate the weight vector Φ*^n^* of the learning network. We then assess how well the *n*-th learning network can predict the actions taken in 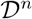. The total reconstruction loss is based on the negative-log-likelihood:

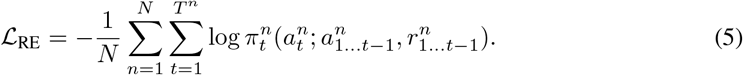

### Disentanglement loss

The second term in the loss function favors disentangled representations by seeking to ensure that each dimension of the latent space corresponds to an independent factor of contribution in the variation across input sequences. This is achieved by maximising the similarity between the empirical encoding distribution (i.e., {**z***^n^*}) and a prior distribution *p*(**z**), taken to be isotropic Gaussian (see the Supplementary Material for details). Maximising this similarity encourages the dimensions of the latent representation to be independent across input sequences. Define 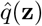 as the empirical distribution of **z***^n^*, and *g*(**z**) as a Gaussian with the same mean and covariance as that of 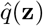. We combine two popular measures of the dissimilarity between the encoding distribution and *p*(**z**):

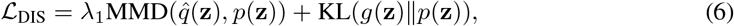

where MMD is the Maximum Mean Discrepancy [Tolstikhin et al., 2017] and KL is the Kullback-Leibler divergence (the Gaussian approximation makes the KL computations tractable and robust) and λ_1_ is a hyper-parameter. We expected the MMD term to suffice; however, we found that combining the two yielded better results.

### Separation loss

Crudely speaking, the disentanglement loss focuses on regularising the encoder; however, to make the latent representation behaviorally interpretable, it is also necessary to regularize the decoder. Gaining insight into the effect of one dimension on the behavioral predictions of the learning network is hard if this is affected by the other dimensions. Therefore, we introduce an additional separation loss designed to discourage interactions.

For simplicity, assume the decision-making task involves only two actions, *C*_1_ and *C*_2_, and that the latent space has two dimensions, *z*_1_ and *z*_2_ (*M* = 2; see Supplementary Material for the general case). As noted before 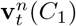 and 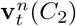 are the logits corresponding to the probability of taking action *C*_1_ and *C*_2_ at trial *t* for input sequence *n*. Denote by 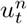 the relative logits of the actions, 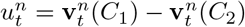. The amount that 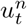 changes by changing the first dimension, *z*_1_ is

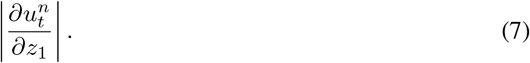

Ideally, the effect of changing *z*_1_ on behavior will be independent from the effect of *z*_2_ on behavior, which will allow us later to interpret *z*_1_ independently from *z*_2_. We capture the amount of interaction (inseparability) between the effect of *z*_1_ and *z*_2_ on changing 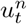 as

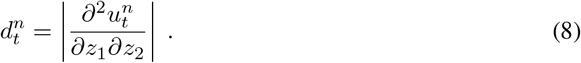

This would be zero if the relative logit comprises *additively separable functions*, i.e., 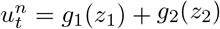 for two functions *g*_1_ and *g*_2_. Even without this, minimizing 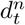 can help make each dimension have a separate effect on behavior. We therefore consider the following loss term,

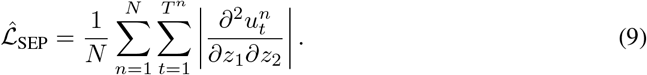

The calculation of the above term is computationally intensive as it requires computing the second-order derivative for each time step and sequence pair (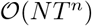 second-order derivative calculations). Instead, we use the following approximation,

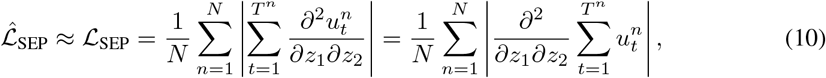

which can be calculated more efficiently (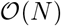 second-order derivative calculations)^1^. We note that the loss is defined using logits instead of the probabilities, since probability predictions are bounded and cannot be separated as 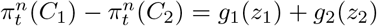 in the general case.

We now define the combined loss function as follows,

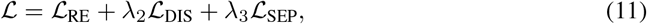

in which λ_2_ and λ_3_ are hyper-parameter.

The model parameters Θ_enc_ and Θ_dec_ were trained based on the above objective function and using gradient descent optimisation method [Kingma and Ba, 2014]. See Supplementary Material for details.

## 4 Results

### synthetic data

To illustrate that our method can learn the underlying ground truth dynamical system, we generated *Q*–learning agents [Watkins, 1989] with various parameter values and simulated their behaviour on a bandit task involving two stochastically-rewarded actions, *C*_1_ and *C*_2_. The actions of the agents and the rewards they received comprise the dataset 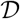. The *Q*–learning agent was specified by two parameters, values for which were drawn randomly (see Supplementary Material for more details). One parameter is the inverse temperature or reward sensitivity, *β*, which determines the propensity to explore, equivalently controlling the impact of receiving a reward from an action on repeating that action in future trials. The other parameter is *κ*, which determines the tendency of an agent to repeat the last taken action in the next trial irrespective of whether it was rewarded [e.g., Lauand Glimcher, 2005]. Values *κ* > 0 favor repeating an action in next trial (perseveration) and values *κ* < 0 favor switching between the actions. We generated *N* = 1500 agents (saving 30% for testing). The test data was used for determining the optimal number of training iterations (early stopping). Each agent selected 150 actions (*T^n^* = 150).

We trained our model using the data (see Figure S3 for the trajectory of the loss function and Figure S4 for the distribution of the latent representations). As shown in Figure 2(a), and as intended, these representations turned out to have an interpretable relationship with the parameters that actually determined behavior: the exploration parameter *β* is mainly related to *z*_2_ and the perseveration parameter *κ* to *z*_1_. This outcome arose because of the 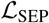 term in the loss function. A similar graph ***before*** introducing the 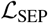 is shown in Figure S1(a), which shows that without this term, *z*_1_ and *z*_2_ have mixed relationships with *κ* and *β*.

**Figure 2:**
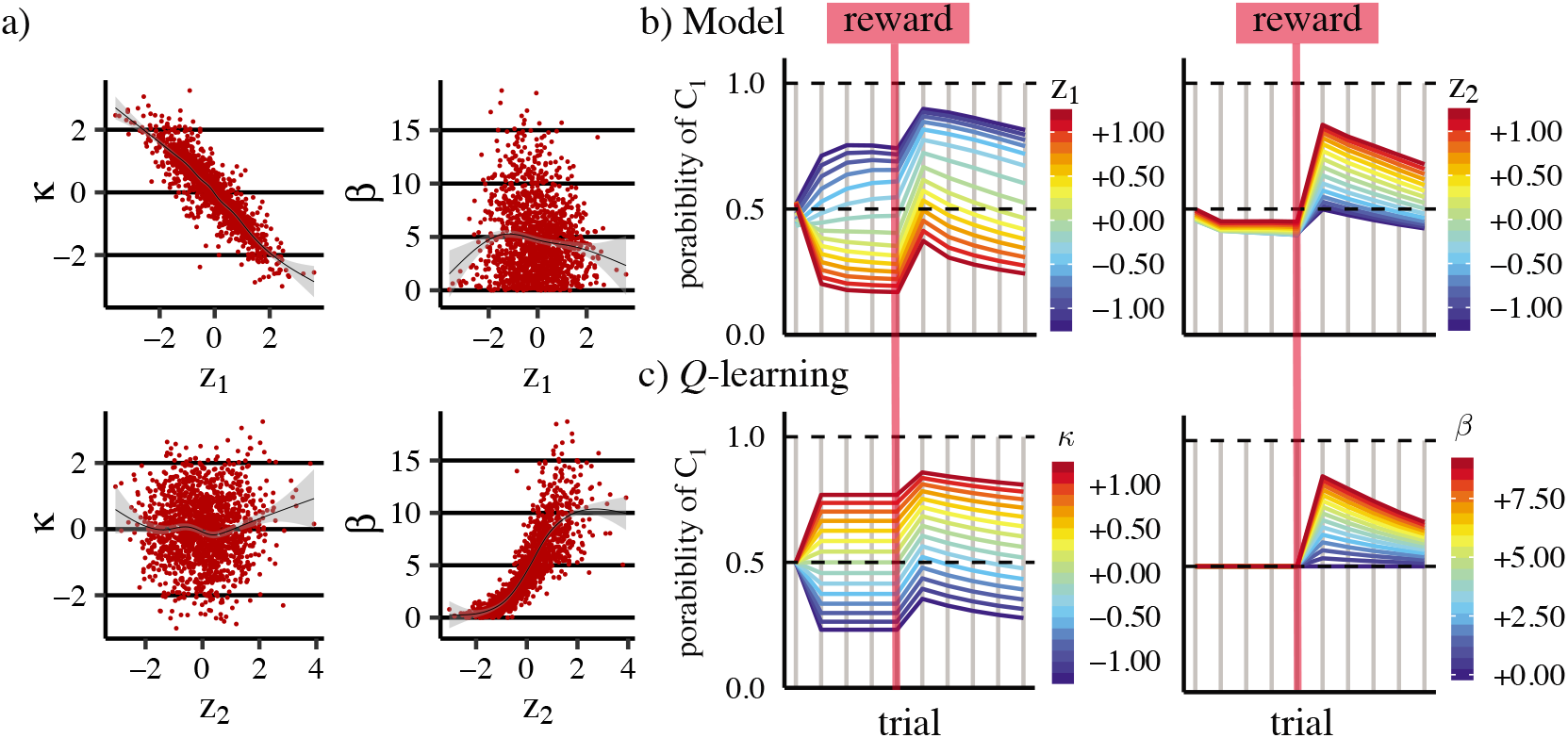
synthetic data. (a) Relationship between the dimensions of the latent representations (*z*_1_, *z*_2_) and the parameters used to generate the data (*κ* and *β*). *z*_1_ captures the perseveration parameter (*κ*) and *z*_2_ captures the rate of exploration (*β*). The smoothed black lines were calculated using method ‘gam’ in R [Wood, 2011] and the shaded area represents confidence intervals. (b) Off-policy simulations of the model for different values of *z*_1_ (left-panel; *z*_2_ = 0) and *z*_2_ (right-panel; *z*_1_ = 0). The plots show the probability of selecting *C*_1_ in each trial when *C*_1_ had actually been chosen on all the previous trials. A single reward is provided, shown by the vertical red line. (c) Model simulations similar to the ones in panel (b) but using the actual *Q*–learning model. In the left panel *β* = 3 and in the right panel *κ* = 0.

We then sought to interpret each dimension of the latent space in behavioral terms. To do this, we used the decoder to generate learning networks corresponding to different values of the latent representations and interpreted these dimensions by analysing the behaviour of the simulated networks on a fixed input sequence. In the fixed input sequence, the agent always performs action *C*_1_ (see Figure S5 for the simulations using both *C*_1_ and *C*_2_ as the previous action), and receives just one non-zero reward at the trial marked by the red vertical line. Using the same sequence for all networks requires running simulations off-policy (i.e., the network predicts the next choice, but does not execute it). This setting provides the same input to the model in all conditions, allowing us to diagnose exactly what affects the output of the model. The simulations are shown in Figure 2(b). Each panel in the figure shows the simulation of the generated network for 10 trials.

In Figure 2(b)-left panel, *z*_2_ is fixed at zero as *z*_1_ is allowed to vary. As the figure shows, by changing the *z*_1_ dimension, the probability of taking the same action (in this case *C*_1_) in the next trial is affected, i.e., high values of *z*_1_ are associated with the high probability of perseveration, and in the low values of *z*_1_ there is a high chance of switching to the other action in the next trial. This is consistent with the correspondence between *z*_1_ and the perseveration parameter *κ*. Note that after receiving the reward, the probability of taking *C*_1_ increases (since *C*_1_ was the previous action and it was rewarded), however, the size of the increment is not affected by changes in *z*_1_, i.e., *z*_1_ is controlling perseveration but not sensitivity of behaviour to reward.

In contrast in Figure 2(b)-right panel, *z*_1_ is fixed at zero as *z*_2_ is in turn allowed to vary. As the panels show, by changing *z*_2_ the probability of repeating an action is not affected, but the sensitivity of action probabilities to the reward is affected. That is, at the higher values of *z*_2_ there is a high probability of repeating the rewarded action (*C*_1_ in this case). Therefore, *z*_1_ and *z*_2_ have separate effects on behaviour corresponding to the perseveration and reward sensitivity. This separation is a consequence of introducing the 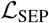 term. In Figure S1(b) we show the same simulations but without introducing the 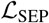 into the loss function, which shows that the effects of *z*_1_ and *z*_2_ on behaviour are not separated (see Figure S2 for how the behaviour of model changes during training). Figure 2(c) shows the same simulations as panel (b) but using the original *Q*–learning model which was used to generate the data (for different values of *β* and *κ*). As the figure shows the model closely reflects the behaviour of *Q*–learning model, as expected.

### bd dataset

This dataset [Dezfouli et al., 2019] comprises behavioural data from 34 patients with depression, 33 with bipolar disorder and 34 matched healthy controls. As in the synthetic data above, subjects performed a bandit task with two stochastically-rewarded actions (*C*_1_ and *C*_2_) (Figure 3(a)). Each subject completed the task 12 times using different reward probabilities for each action. The dataset thus contains *N* = 12 (sequences) × 101 (participants) = 1212, which we used for training the model. Out of the 12 sequences of each subject, 8 were used for training and 4 for testing to determine the optimal number of training iterations (see Figure S7 for the training curves and Supplementary Material for more details).

**Figure 3:**
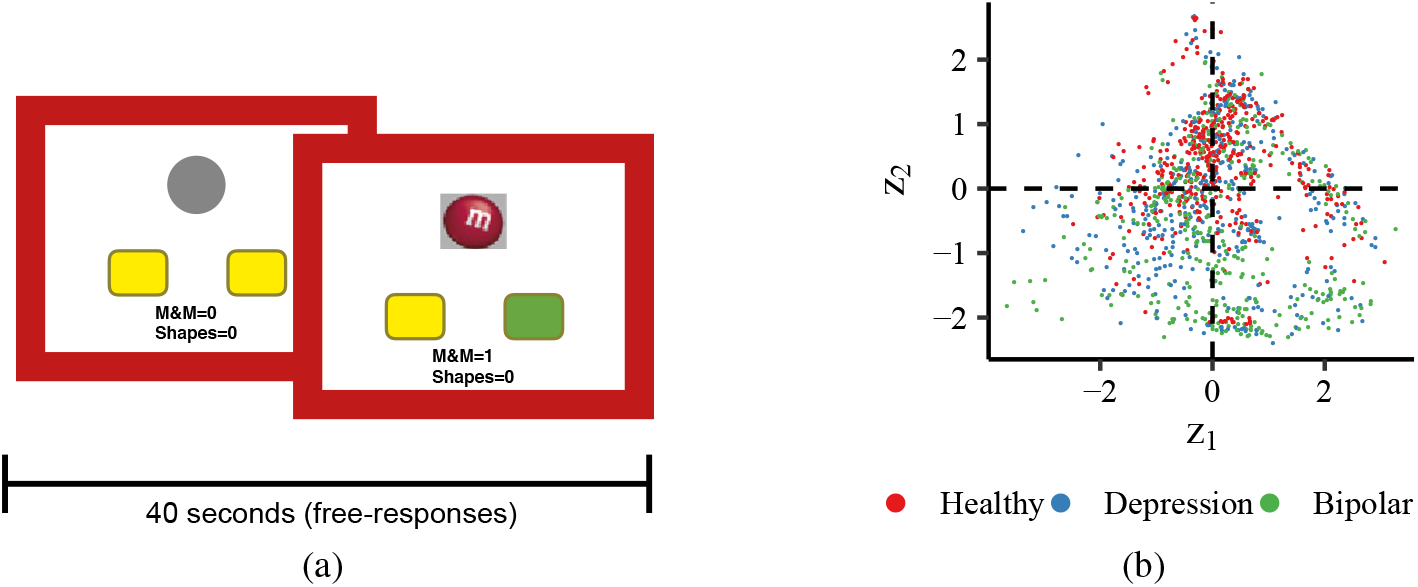
bd dataset. (a) The decision-making task. On each trial, subjects pressed a left (*C*_1_) or right (*C*_2_) key and had a chance of getting a reward (M&M chocolate or BBQ shapes). The task lasted for 40 seconds and the responses were self-paced. Each participant completed the task 12 times with a 12-second delay between them and different probabilities of reward. (b) Distribution of **z** values for each group. Each dot represents an input sequence.

We considered a two-dimensional latent space **z** = { *z*_1_, *z*_2_}; the resulting coordinates for all sequences are shown in Figure 3(b). We made two predictions about **z**: first, we expected that the latent representations for the sequences for a single subject should be mutually similar as they come from the same decision-making system. We therefore compared the mean pairwise distances separating the latent representations within and between subjects (see Supplementary Material for details). Figure 4(a) shows that this prediction is indeed correct (*p* < 0.001 using Wilcoxon rank sum test).

**Figure 4:**
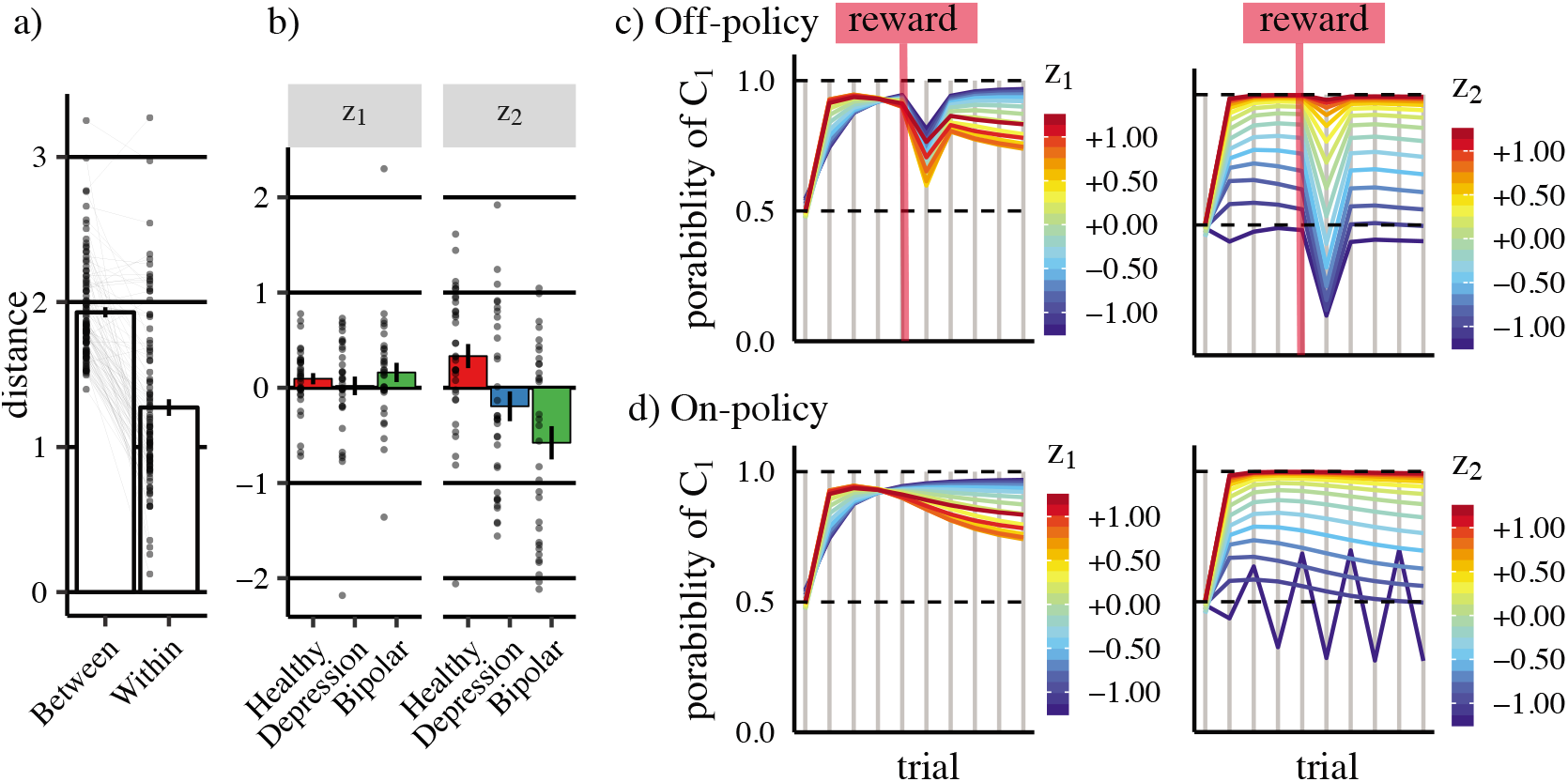
bd dataset. (a) The distances between the latent representations for single subjects (Within) and between the subjects (Between). Each dot represents a subject; the bars show the means and the error-bars, 1SEM. (b) The mean of the latent representations for each subject across the dimensions of the latent space. Each dot represents a subject and bars and error-bars show means and 1SEM. (c) Off-policy simulations of the model for different values of *z*_1_ (left-panel; *z*_2_ = 0) and *z*_2_ (right-panel; *z*_1_ = 0). The plots show the probability of selecting *C*_1_ in each trial when *C*_1_ had actually been chosen on all the previous trials. A single reward is provided, shown by the vertical red line. (d) On-policy simulations of the model. The actions are selected by the model based on which action has the higher probability of being taken (the first action was set to *C*_1_).

Second, we expected that the decision-making differences between the three groups to be reflected in their latent representations. Figure 4(b) shows the mean latent representations for the subjects organized by group. Although the groups differ along the *z*_2_ dimension (*p* < 0.001 comparing bipolar and healthy groups along *z*_2_ and using independent t-test, and *p* < 0.05 comparing depression and healthy groups); the *z*_1_ dimension is evidently capturing variations that are unrelated to the existing diagnostic categories (*p* > 0.1).

These results also highlight the substantial heterogeneity of the population. Some bipolar patients with high *z*_2_ are apparently more behaviorally similar to average depressed patients or healthy controls than to the average bipolar patients. We would not have been able to extract this information by fitting a single RNN to each whole group (as was done in previous work).

We then followed the same scheme as for the synthetic data to provide an interpretation for the dimensions of latent space. The results of off-policy simulations are shown in Figure 4(c) (see Figure S6 for simulations using both actions). First, both left and right panels show that, after receiving the reward from an action, subjects show a tendency to *switch* to the other action. This is inconsistent with predictions of conventional *Q*–learning, but was also found to be evident in model agnostic analyses [Dezfouli et al., 2019] (where it is also reconciled with gaining rather than losing reward). The current model is able to capture this since it uses RNNs for representing decision-making processes, which do not make such an assumption.

Second, as Figure 4(c)-left shows, the *z*_1_ dimension – which is not different between the groups – is mostly related to reward sensitivity, as it controls (albeit rather weakly) the action probabilities after receiving the reward. On the other hand, the *z*_2_ dimension – which is significantly different between the groups – is more involved in perseveration/switching behaviour between the actions, i.e., it controls the probability of staying on the same action. For low *z*_2_, the probability of repeating the previous taken (in this case *C*_1_) is below 0.5, implying that the subjects in this part of the latent space tend to switch. In order to confirm this, we simulated the model on-policy – in which the actions are selected by the model – for different values of *z*_1_ and *z*_2_. As the results in Figure 4(d) show, for low *z*_2_, the model indeed oscillates between the two actions. The *z*_2_ dimension is significantly lower in the bipolar group than healthy controls, which is consistent with the previous report indicating an oscillatory behavioural characteristic in this group [Dezfouli et al., 2019].

Finally, panel (c)-right shows that reward sensitivity and perseveration are not completely independent, i.e., when *z*_2_ is low, favoring switching, the effect of reward on probabilities is also more significant, implying that these two traits covary.

In summary, the model was able capture behavioural properties which are not consistent with cognitive models such as *Q*–learning. It was also able to capture individual differences which could not be extracted by fitting a single RNN to the whole group.

## 5 Related work

There is a wealth of work using RNNs as models of decision-making, for unsupervised dimension reduction of dynamical systems, and for sequence-to-sequence mappings. For the first, a key focus has been learning-to-learn [e.g., Wang et al., 2016, Song et al., 2017] – i.e., creating RNNs that can themselves learn to solve a battery of tasks. However, these generative models have not been coupled with recognition, for instance to capture individual differences. Techniques based on autoencoders have explored low-dimensional latent spaces for modelling neural activity trajectories in the brain [e.g., Pandarinath et al., 2018]. However, like influential sequence-sequence models for translation [Bowman et al., 2015], these focus on building an open loop account for the *state* of the brain within a trajectory, or the *state* characterizing a particular input sentence, whereas we focus on the *trait* characteristic of a closed-loop controller that has to interact with an external environment itself [see also Fox et al., 2011, Johnson et al., 2016, as examples of other approaches]. Our combination of an autoencoder and hyper-network [Ha et al., 2016, Karaletsos et al., 2018] is, to our knowledge, novel, and might find other applications in the analysis of other dynamical systems such as time-series data.

## 6 Conclusion and discussion

We proposed a flexible autoencoder-based framework for modelling individual differences in decisionmaking tasks. The autoencoder maps the sequence of actions and rewards taken and received by a subject to a position in a low-dimensional latent space which is decoded to determine the weights of a ‘learning network’ RNN that characterizes how that subject behaves as a plastic decision-maker. The latent space was disentangled by adding a specific component to the loss function.

The model can be extended in various directions. One is to pin down (part of) the learning network as a conventional, parameterized, RL agent, to facilitate further interpretation of the latent dimensions. Another is to include tasks with non-trivial (Markovian or non-Markovian) state information partially signalled by stimuli.

## Acknowledgments

We are grateful to Bernard W. Balleine for sharing BD dataset with us.

## Supplementary Material

### S1 The model

#### S1.1 Model architecture

The recurrent neural network in the encoder network consists of *N*_enc_ bidirectional LSTM cells [Hochreiter and Schmidhuber, 1997, Schuster and Paliwal, 1997]. The final output vectors of the RNN (backward and forward) are concatenated (which will be of dimensionality 2*N*_enc_) and fed into four feed-forward fully-connected layers, with *N*_enc_, *N*_enc_, 10, and 2 neurons respectively. The last layer corresponds to the latent representations (in our case, with 2 neurons). The activation function for the layers are Rectified linear units [ReLU; Nair and Hinton, 2010], *ReLU, soft-plus* [Dugas et al.,2001] and linear respectively. The decoder network is composed of four fully-connected feed-forward layers with 100,100,100, 69 neurons respectively and with *tanh* activation function in all the layers. The outputs of the last layer (69 outputs) are used as the weights of the GRU-based learning network and the mapping *W^n^* from this network to the policy. The GRU network has 3 cells, which requires 63 weights, and *W^n^* has 3 (GRU outputs) ×2 (actions) elements.

The learning network is small to avoid its being able to predict the input signal without relying on the latent representations, a phenomenon known as *posterior collapse* in variational autoencoders [e.g., Bowman et al., 2015, Kingma et al., 2016]. To assess the extent of posterior collapse, we calculated a *random* reconstruction error during training by assessing how well a network generated based on the latent representation of one sequence could predict the actions of another. If the learning network is weak enough that it cannot ignore the latent representation, then its performance should be compromised by this shuffling. Figures S3 and S7 show that the random reconstruction loss is indeed higher than the training loss for both the synthetic and BD data, which indicates that posterior collapse has been successfully inhibited.

#### S1.2 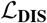 term

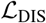 is composed of two components: an MMD term and a KL term,

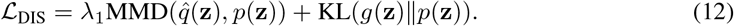

For the calculation of the MMD term, a Gaussian Radial basis function (RBF) kernel was used to turn the samples **z***^n^* into a smooth distribution. The kernel hyperparameters that maximized the MMD were used.^2^

In principle, the MMD term should suffice to force the latent distribution to follow the prior distribution *p*(**z**). However, there are other measures of discrepancy between distributions, and the possibility of combining various such. We found it best to include contributions from the KL divergence between *p*(**z**) and a Gaussian match (*g*(**z**)) of the mean (**m_z_**) and covariance (Cov**_z_**) of 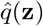.

Theoretically, combining the two losses in fact makes sense from a geometric standpoint. Under some assumptions, the MMD term can be related to integral probability metrics and then Kantorovich-Rubinstein formula [Villani, 2009, Particular case 5.16]: it is thus an optimal transport dissimilarity between the supports of 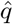 and *p* (the data manifold). On the other hand, since *g* and *p* are Gaussian, it is also a known result that the KL divergence between two Gaussians equals a sum of the LogDet divergence between their covariance matrices [Kulis et al., 2009] and a Mahalanobis divergence between their averages [Amari., 1985]: it thus defines an information geometric distortion between their parameters [Amari and Nagaoka, 2000] (the statistical manifold).

#### S1.3 Training the model

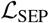 can itself induce a local optimum form of collapse associated with ignoring one of the latent dimensions. If, for instance, 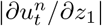 is uniformly small, then 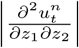, and thus 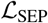 will be small. This can persist as the gradients of the weights that pass through the ignored latent dimension will continue to remain small throughout the training process and therefore the weights will not get updated.

Thus, we trained the model in stages. In stage 1, we started by setting λ_3_ = 0 (eliminating 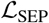) and chose the optimal number of training iterations using cross-validation (early-stopping). Then, in stage 2, we introduced 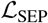 term using a small λ_3_ (λ_3_ ≤ 1), whist ensuring that the reconstruction loss was not materially affected, thus avoiding collapse.

All the gradients were calculated using automatic differentiation and the models were optimised using Adam optimiser [Kingma and Ba, 2014]. In the stage 1 of training, all the parameters (Θ_enc_ and Θ_dec_) were optimised together. In the stage 2 of training (in which 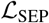 was introduced), Θ_enc_ and Θ_dec_ were optimised iteratively. That is, Θ_dec_ was optimised for 50 times and then Θ_enc_ was optimised for 200 times, and then this loop was iterated.

Since some subjects might have a bias towards one of the actions (*C*_1_ or *C*_2_), we randomly counterbalanced *C*_1_ and *C*_2_ between the encoder and decoder in order to prevent the latent representations from being affected by such biases.

We used an Accelerator Cluster for running the experiments (with NVidia Tesla P100 (SXM2)).

#### S1.4 Model parameters

For bd data we set *N*_enc_ = 20. For synthetic data, we observed that *N*_enc_ = 20 did not fit the behaviour adequately (i.e., reconstruction error was still decreasing after >60000 training iterations without increasing the test reconstruction error), and therefore we used a more powerful encoder by choosing *N*_enc_ = 50 for synthetic experiment.

In stage 1 of the optimisation (i.e., loss without 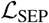 term – see above) the hyper-parameters were λ_1_ = 50, λ_2_ = 1 and λ_3_ = 0. For the second stage of the optimisation the hyper-parameters were λ_1_ = 50, λ_2_ = 1, λ_3_= 1 for synthetic experiment. For BD experiment we found that λ_3_ = 1 at stage 2 of training causes an increase in the reconstruction loss during optimisation, which was then avoided by decreasing λ_3_ to 0.1. The other two parameters were also decreased to λ_1_ =2, λ_2_ = 0.5. In this settings the 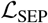 decreased without materially increasing 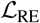 during stage 2 optimisation.

#### S1.5 On-policy simulations

For the on-policy simulations, the actions were selected by model based on the probabilities predicted by the model. The first action that was fed to the model was *C*_1_ and the remaining actions were selected by the model.

### S2 synthetic data

For generating synthetic data, the agents learned and selected actions as follows. At each time step *t*, after agent *n* selected action 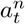 and received reward 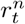, the *Q*-value of action 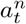 was updated according to 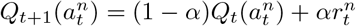. *α* is the learning rate, and was fixed to 0.2 for all the agents. The action in the next trial was selected according to the following probability,

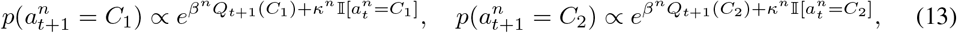

in which 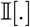 is the indicator function.

The perseveration parameter for agent n was drawn randomly from a Gaussian distribution,

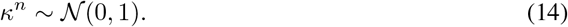

The inverse temperature parameter was selected randomly according to,

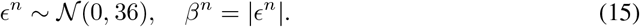

We generated 1500 agents, each of which selected 150 actions. 30% of agents were used for testing. The actions paid off probabilistically as either {*p*(*r* = 1|*a* = *C*_1_) = 0.1, *p*(*r* = 1|*a* = *C*_2_) = 0.5}, or {*p*(*r* = 1|*a* = *C*_2_) = 0.1, *p*(*r* = 1|*a* = *C*_1_) = 0.5}, counterbalanced randomly across the agents.

Since the latent space has only two dimensions, we were able to directly visualize/report one-to-one relationships between each latent variable and each factor of variation in the data (Figures 2, S1). We also calculated the disentanglement metrics reported in disentanglement_lib [Locatello et al., 2018] based on the results in the synthetic dataset obtained by including the separation loss, and without including the separation loss, which showed that the separation loss improved disentanglement.

### S3 bd dataset

In this dataset, each subject completed the task 12 times (12 blocks), so each generated 12 input sequences. Of those sequences, a randomly selected 8 were used for training the model and the remaining 4 for testing. On average, participants completed 109.45, 114.91, 102.79 trials per block in healthy, depression, and bipolar groups respectively.

To generate Figure 4(a), we calculated the mean pairwise distance between the latent representations of each subject *i* 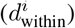, and the pairwise distance between the latent representations of each subject i to the other subjects 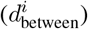. For calculating 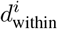, note that there are 12 latent representations for each subject. For each of these representations, we calculated the total of 66 non-trivial, unique, distances arising from the representation of input sequence *i* to that of the input sequence *j* (∀*j* > *i*). Denote by **z***^ik^* the latent representation for the *k*th input sequence of subject *i*,

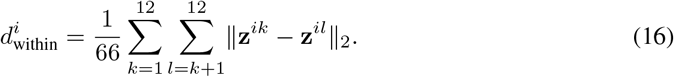

For calculating 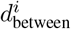, we calculated the distance between latent representations of subject *i* and each subject *j* (*i* ≠ *j*),

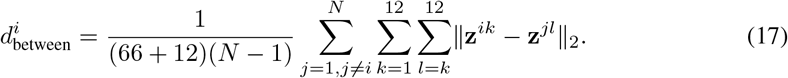

Note that there are 66 + 12 pair of distances between each two subjects, as the distance is symmetric.

### S4 Generalized separation loss

In Section 3 we introduced the separation loss for the special case of only two actions and two latent variables. In this section we extend this notion to address a more general setting. Recall that 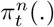 was the vector of probabilities that the learning network assigns to the actions (at trial *t* and for the subject *n*). Let 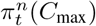 be the maximum value of these action probabilities, and let 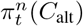 be the second highest of these action probability. Define

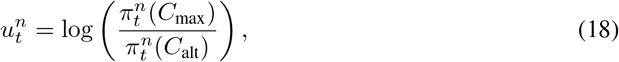

which basically measures the learning network’s confidence in its preferred action (compared to the “alternative” second best action). Note that Eq. 18 is consistent with the definition that we had for 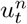 in Section 3 since for the special case of two actions we have

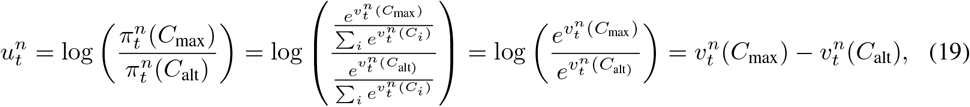

where *C*_max_ and *C*_alt_ are the actions corresponding to 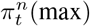 and 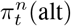 respectively. Ideally, the effect of different *z_i_*’s on changing the behaviour (i.e., the preferred action) should be independent of each other. The following notion, which is analogous to Eq. 8, captures the pairwise interactions between *z_i_* and *z_j_* on changing the preferred action

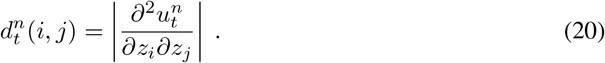

In order to be able to interpret each latent variable independently, we minimize these pairwise interactions over all the choices of *z_i_* and *z_j_*. In particular, we aim at minimizing the following loss function

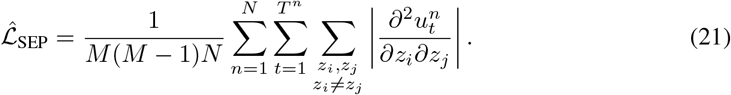

where *M* was the dimensionality of the latent space. Note that for the special case of *M* = 2, Eq. 21 simply recovers the definition that we already had in Eq. 9

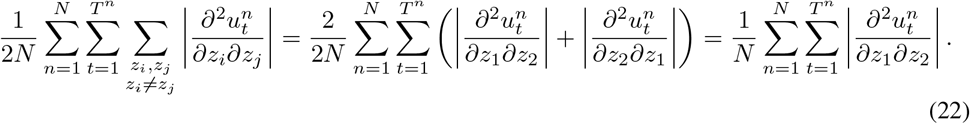

---

**Figure S1:**
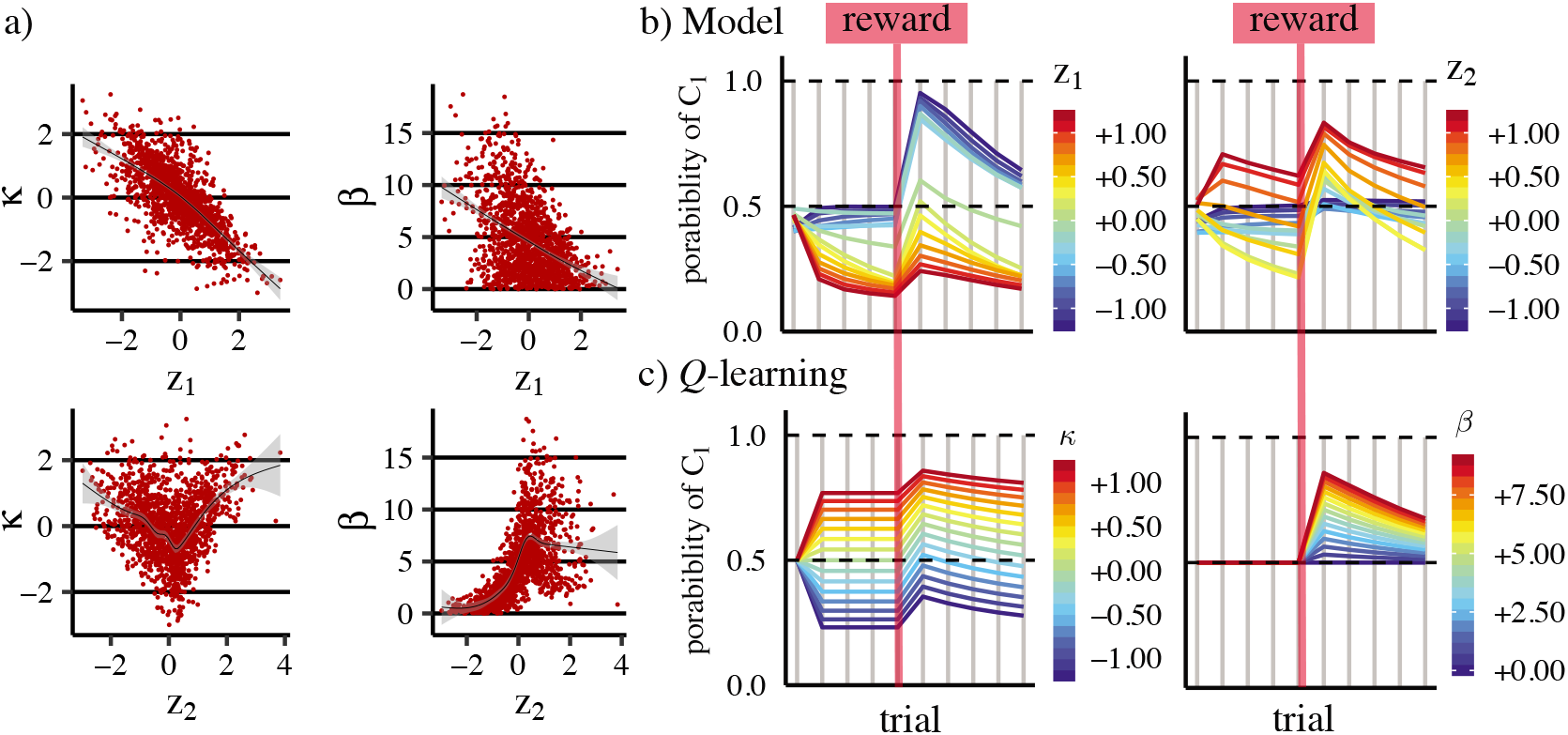
This figure is similar to Figure 2 but *without* optimising the loss function with 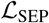 term. (a) Relationship between the dimensions of the latent representations (*z*_1_, *z*_2_) and the parameters used to generate the data (*κ* and *β*). The black lines were calculated using method ‘gam’ in R [Wood, 2011] and the shaded area shows confidence intervals. (b) Off-policy simulations of the model for different values of *z*_1_ (left-panel; *z*_2_ = 0) and *z*_2_ (right-panel; *z*_1_ = 0). The plots show the probability of selecting *C*_1_ in each trial when *C*_1_ had actually been chosen on all the previous trials. A single reward is provided, shown by the vertical red line. (c) Model simulations similar to the ones in panel but using the actual *Q*–learning model. In the left panel *β* = 3 and in the right panel *κ* = 0.

**Figure S2:**
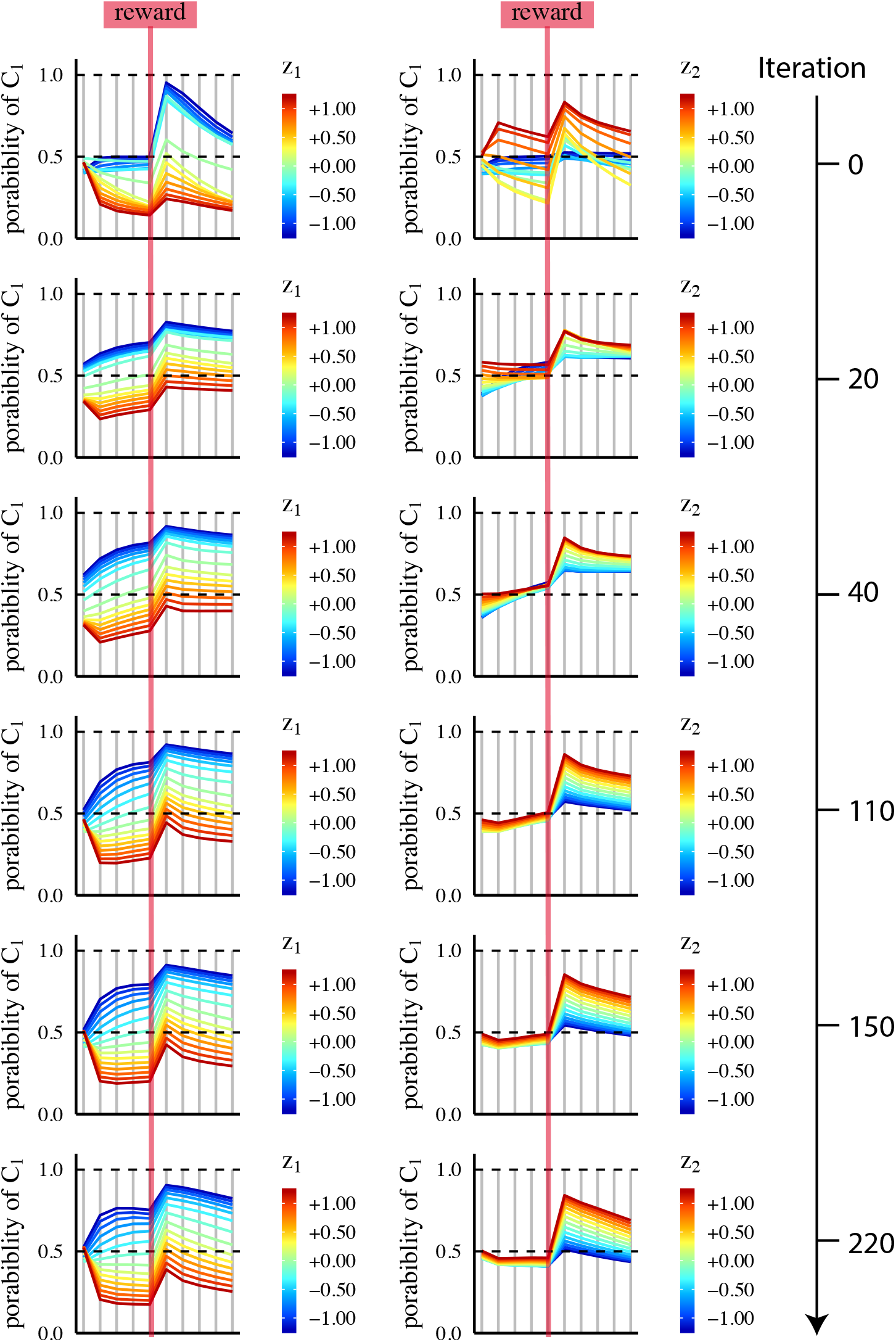
synthetic dataset. Off-policy simulations during model training, which shows how the effect of *z*_1_ and *z*_2_ on behaviour are separated after the introduction of 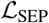 term. Note that iteration 0 is the beginning of the introduction of 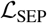 term in the loss function. Each iteration consists of 50 updates of decoder parameters and 200 updates of encoder parameters. Note that this is only stage 2 of the training and after the introduction of 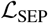 term into the loss function.

**Figure S3:**
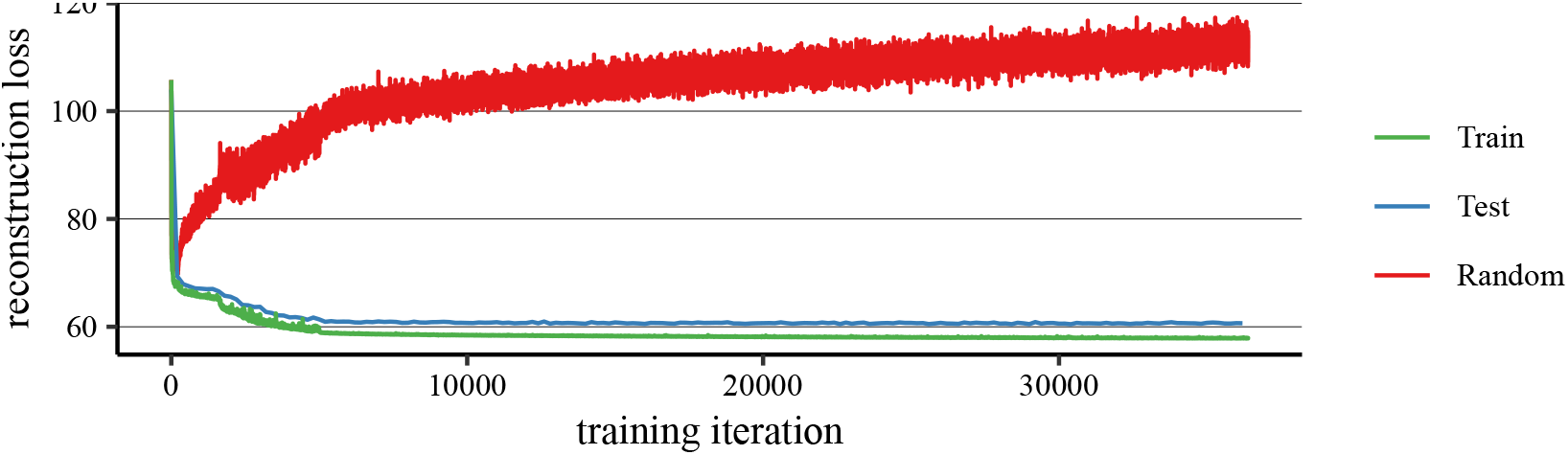
synthetic data. Training, test, and random reconstruction loss. Random reconstruction loss quantifies the specificity of the latent representations to each input sequence, allowing us to detect posterior collapse. To calculate this loss, the learning network for a sequence is generated based on the encoded latent representation of a radomly-selected *different* sequence. If the latent representations are specific to the input sequences we expect this random reconstruction loss to increase by training, which is the case as shown in the graph. Note that the graph only shows stage 1 of the training. See text for more description.

**Figure S4:**
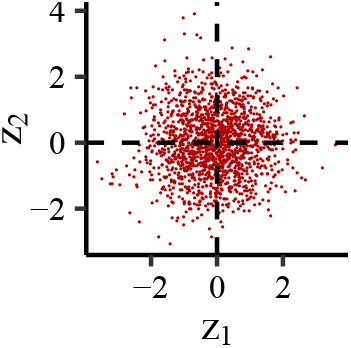
synthetic data. Distribution of *z* values.

**Figure S5:**
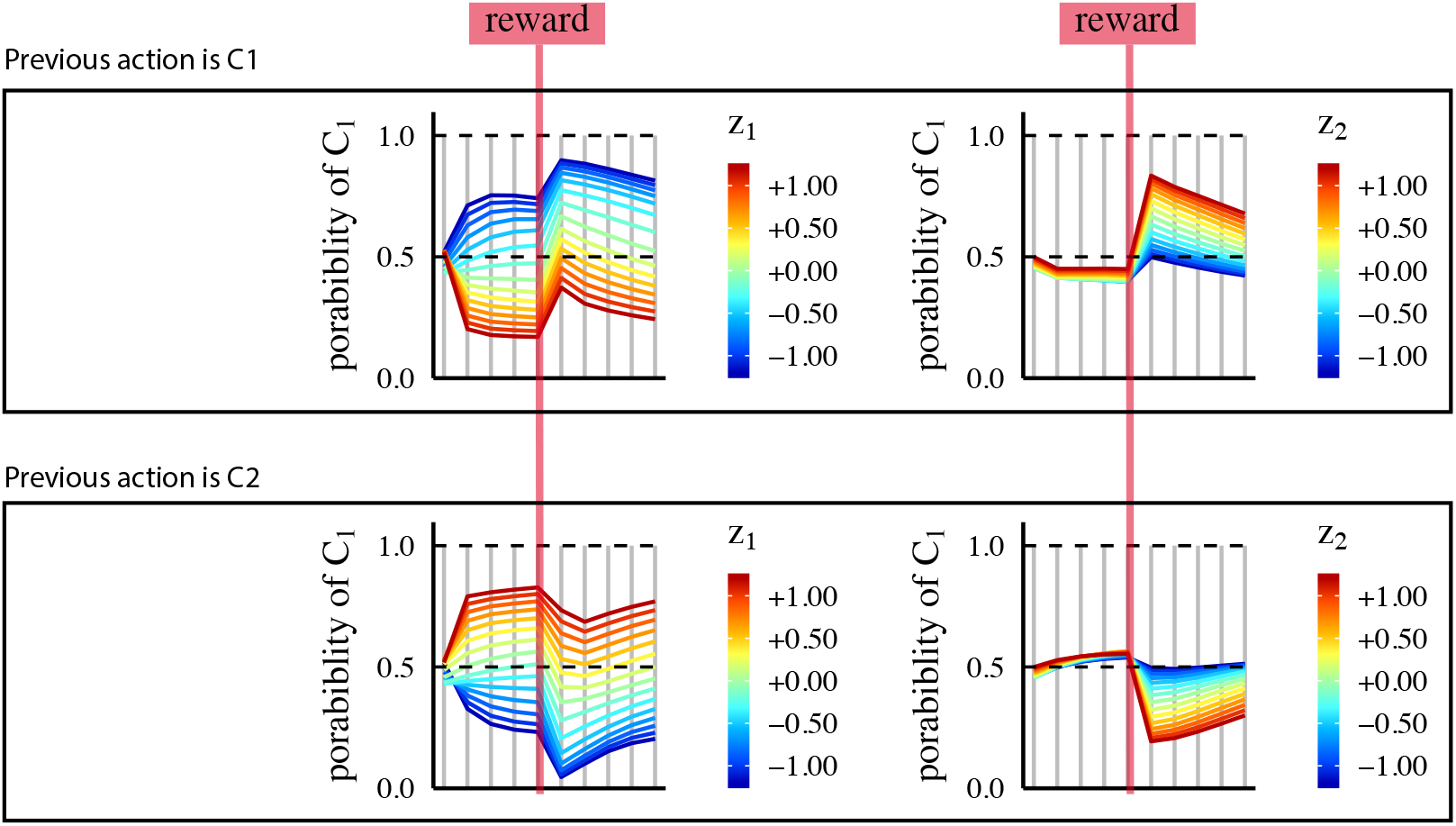
synthetic data. Off-policy simulations of the model. The top panel is similar to Figure 2(b). The bottom panel is similar to top panel, but the action fed to the model as the previous action was *C*_2_ (instead of *C*_1_ which was fed to the model in the top panel).

**Figure S6:**
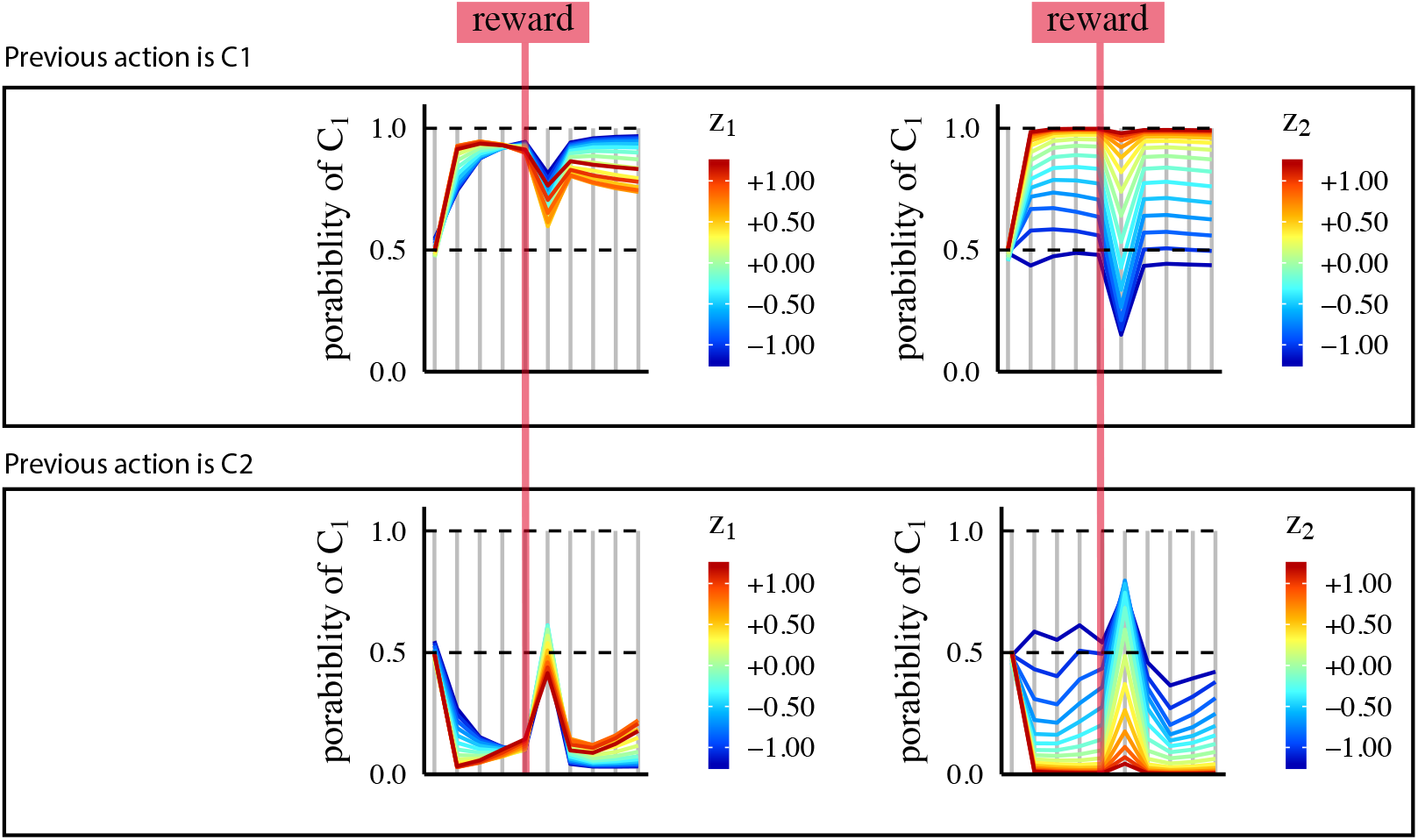
bd data. Off-policy simulations of the model. The top panel is similar to Figure 4(c). The bottom panel is similar to the top panel, but action *C*_2_ was fed to the model as the previous action (instead of *C*_1_ which was fed the model in the top panel).

**Figure S7:**
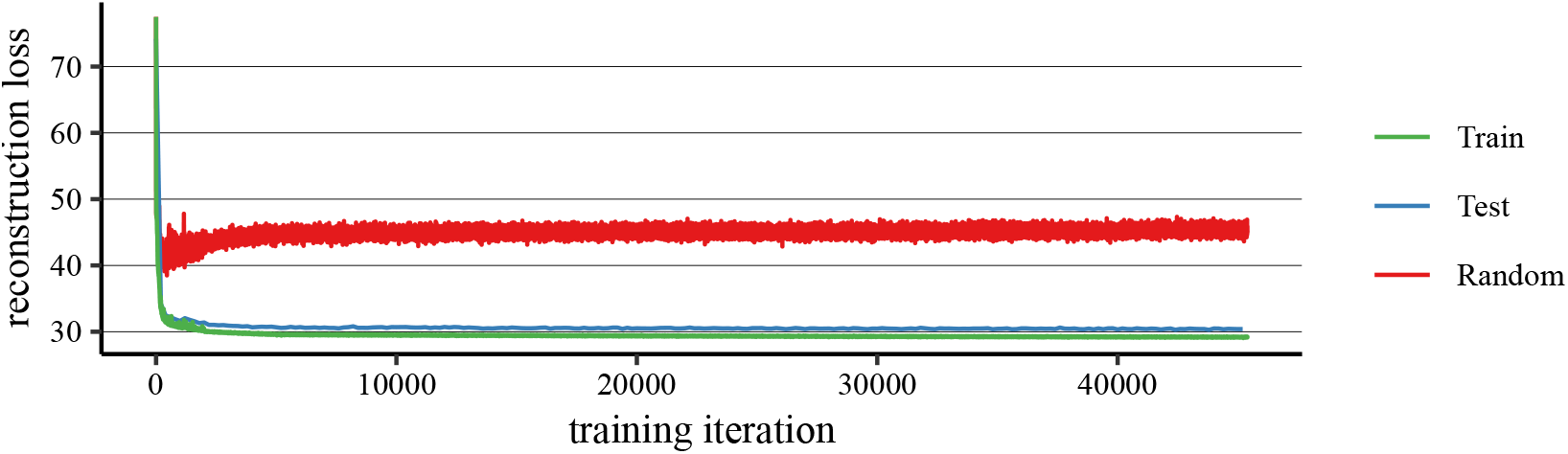
bd data. Training, test, and random reconstruction loss. Random reconstruction loss quantifies the specificity of the latent representations to each input sequence, allowing us to detect posterior collapse. To calculate this loss, the learning network for a sequence is generated based on the encoded latent representation of a radomly-selected *different* sequence. If the latent representations are specific to the input sequences we expect this random reconstruction loss to increase by training, which is the case as shown in the graph. Note that the graph only shows stage 1 of the training. See text for more description.

## Footnotes

1 We use the automatic differentiation in Tensorflow [Abadi et al., 2016].

2 Employing the implementation: https://github.com/tensorflow/models/blob/master/research/domain_adaptation/domain_separation/losses.py

## References

John B Carroll and Scott E Maxwell. Individual differences in cognitive abilities. Annual review of psychology, 30(1):603–640, 1979.

Michael J Frank, Bradley B Doll, Jen Oas-Terpstra, and Francisco Moreno. Prefrontal and striatal dopaminergic genes predict individual differences in exploration and exploitation. Nature neuroscience, 12(8):1062, 2009.

Hanneke E M den Ouden, Nathaniel D Daw, Guillén Fernandez, Joris A Elshout, Mark Rijpkema, Martine Hoogman, Barbara Franke, and Roshan Cools. Dissociable effects of dopamine and serotonin on reversal learning. Neuron, 80(4):1090–1100, 2013.

Daniel J Navarro, Thomas L Griffiths, Mark Steyvers, and Michael D Lee. Modeling individual differences using Dirichlet processes. Journal of mathematical Psychology, 50(2):101–122, 2006.

Jerome R Busemeyer and Julie C Stout. A contribution of cognitive decision models to clinical assessment: decomposing performance on the Bechara gambling task. Psychological assessment, 14(3):253, 2002.

Nathaniel D Daw. Trial-by-trial data analysis using computational models. In Mauricio R. Delgado, Elizabeth A. Phelps, and Trevor W. Robbins, editors, Decision Making, Affect, and Learning. Oxford University Press, 2011.

Eldad Yechiam, Jerome R Busemeyer, Julie C Stout, and Antoine Bechara. Using cognitive models to map relations between neuropsychological disorders and human decision-making deficits. Psychological science, 16(12):973–8, dec 2005.

Amir Dezfouli, Kristi Griffiths, Fabio Ramos, Peter Dayan, and Bernard W Balleine. Models that learn how humans learn: the case of decision-making and its disorders. PLoS computational biology, 15(6):e1006903, 2019.

Amir Dezfouli, Richard W Morris, Fabio Ramos, Peter Dayan, and Benrard W Balleine. Integrated accounts of behavioral and neuroimaging data using flexible recurrent neural network models. In Advances in Neural Information Processing Systems (Neurips), 2018.

Guangyu Robert Yang, Madhura R Joglekar, H Francis Song, William T Newsome, and Xiao-Jing Wang. Task representations in neural networks trained to perform many cognitive tasks. Nature Neuroscience, 22(2):297–306, 2019.

Hava T Siegelmann and Eduardo D Sontag. On the computational power of neural nets. Journal of computer and system sciences, 50(1):132–150, 1995.

David E Rumelhart, Geoffrey E Hinton, and Ronald J Williams. Learning internal representations by error propagation. Technical report, California Univ San Diego La Jolla Inst for Cognitive Science, 1985.

Ilya Tolstikhin, Olivier Bousquet, Sylvain Gelly, and Bernhard Schoelkopf. Wasserstein auto-encoders. arXiv preprint arXiv:1711.01558, 2017.

David Ha, Andrew Dai, and Quoc V Le. Hypernetworks. arXiv preprint arXiv:1609.09106, 2016.

Theofanis Karaletsos, Peter Dayan, and Zoubin Ghahramani. Probabilistic meta-representations of neural networks. arXiv preprint arXiv:1810.00555, 2018.

Kyunghyun Cho, Bart Van Merriënboer, Caglar Gulcehre, Dzmitry Bahdanau, Fethi Bougares, Holger Schwenk, and Yoshua Bengio. Learning phrase representations using RNN encoder-decoder for statistical machine translation. arXiv preprint arXiv:1406.1078, 2014.

Martín Abadi, Ashish Agarwal, Paul Barham, Eugene Brevdo, Zhifeng Chen, Craig Citro, Greg S Corrado, Andy Davis, Jeffrey Dean, Matthieu Devin, and Others. Tensorflow: Large-scale machine learning on heterogeneous distributed systems. arXiv preprint arXiv:1603.04467, 2016.

Diederik P. Kingma and Jimmy Ba. Adam: A Method for Stochastic Optimization. arXiv preprint arXiv:1412.6980, 2014.

C.J.C.H. Watkins. Learning from Delayed Rewards. Ph.D. thesis, Cambridge University, 1989.

Brian Lau and Paul W Glimcher. Dynamic response-by-response models of matching behavior in rhesus monkeys. Journal of the experimental analysis of behavior, 84(3):555–79, 2005.

Simon N Wood. Fast stable restricted maximum likelihood and marginal likelihood estimation of semiparametric generalized linear models. Journal of the Royal Statistical Society: Series B (Statistical Methodology), 73(1):3–36, 2011.

Jane X Wang, Zeb Kurth-Nelson, Dhruva Tirumala, Hubert Soyer, Joel Z Leibo, Remi Munos, Charles Blundell, Dharshan Kumaran, and Matthew M Botvinick. Learning to reinforcement learn. arXiv preprint arXiv:1611.05763, 2016.

H. Francis Song, Guangyu R. Yang, and Xiao Jing Wang. Reward-based training of recurrent neural networks for cognitive and value-based tasks. eLife, 6:1–24, 2017.

Chethan Pandarinath, Daniel J O’Shea, Jasmine Collins, Rafal Jozefowicz, Sergey D Stavisky, Jonathan C Kao, Eric M Trautmann, Matthew T Kaufman, Stephen I Ryu, Leigh R Hochberg, and Others. Inferring single-trial neural population dynamics using sequential auto-encoders. Nature methods, page 1, 2018.

Samuel R Bowman, Luke Vilnis, Oriol Vinyals, Andrew M Dai, Rafal Jozefowicz, and Samy Bengio. Generating sentences from a continuous space. arXiv preprint arXiv:1511.06349, 2015.

Emily B Fox, Erik B Sudderth, Michael I Jordan, Alan S Willsky, et al. A sticky hdp-hmm with application to speaker diarization. The Annals of Applied Statistics, 5(2A):1020–1056, 2011.

Matthew J Johnson, David K Duvenaud, Alex Wiltschko, Ryan P Adams, and Sandeep R Datta. Composing graphical models with neural networks for structured representations and fast inference. In Advances in neural information processing systems, pages 2946–2954, 2016.

## References

Sepp Hochreiter and Jürgen Schmidhuber. Long short-term memory. Neural computation, 9(8): 1735–1780, 1997.

Mike Schuster and Kuldip K Paliwal. Bidirectional recurrent neural networks. IEEE Transactions on Signal Processing, 45(11):2673–2681, 1997.

Vinod Nair and Geoffrey E Hinton. Rectified linear units improve restricted boltzmann machines. In Proceedings of the 27th international conference on machine learning (ICML-10), pages 807–814, 2010.

Charles Dugas, Yoshua Bengio, François Bélisle, Claude Nadeau, and René Garcia. Incorporating second-order functional knowledge for better option pricing. In Advances in neural information processing systems, pages 472–478, 2001.

Durk P Kingma, Tim Salimans, Rafal Jozefowicz, Xi Chen, Ilya Sutskever, and Max Welling. Improved variational inference with inverse autoregressive flow. In Advances in neural information processing systems, pages 4743–4751, 2016.

C Villani. Optimal transport, old and new. Springer, 2009.

B Kulis, M.-A. Sustik, and I.-S. Dhillon. Low-rank Kernel learning with Bregman matrix divergences. 10:341–376, 2009.

S.-I. Amari. Differential-Geometrical Methods in Statistics. Springer, 1985.

S.-I. Amari and H Nagaoka. Methods of Information Geometry. Oxford University Press, 2000.

Francesco Locatello, Stefan Bauer, Mario Lucic, Sylvain Gelly, Bernhard Schölkopf, and Olivier Bachem. Challenging common assumptions in the unsupervised learning of disentangled representations. arXiv preprint arXiv:1811.12359, 2018.

